# Distinct Disinhibitory Circuits Link Short-Term Adaptation to Familiarity and Reward Learning in Visual Cortex

**DOI:** 10.64898/2026.03.24.713929

**Authors:** A.J. Hinojosa, S.E. Dominiak, Ye. Kosiachkin, L. Lagnado

**Affiliations:** Sussex Neuroscience, School of Life Sciences, University of Sussex, Brighton BN1 9QG, United Kingdom

## Abstract

Sensory cortices filter repeated inputs through rapid adaptation over seconds and experience-driven learning over days. Although these forms of plasticity occur simultaneously, it is not known how they interact within cortical circuits. We combined two-photon calcium imaging, data-driven circuit modelling and optogenetics to investigate how short-term adaptation in layer 2/3 of mouse V1 is shaped by stimulus familiarity and reward association. Habituation reduced the fraction of pyramidal cells responsive to a visual stimulus, whereas reward association maintained overall responsivity. In contrast, both forms of learning shifted pyramidal cell adaptation from depression toward sensitization, but through distinct circuit mechanisms. Habituation reduced disinhibition through the VIP→SST→PC pathway by weakening feedback activation of VIPs and VIP→SST connections. Reward association counteracted this effect by *increasing* disinhibition through the SST→PV→PC pathway, strengthening SST→PV connections while reducing SST→PC inputs. Despite engaging distinct disinhibitory circuits and producing divergent effects on pyramidal cell responsivity, both forms of learning converged on a reduced PV:SST input ratio to pyramidal cells, thereby biasing V1 toward sensitizing adaptation. These results identify changes in cortical circuits that link the plasticity of fast adaptation to simple forms of learning.

## Introduction

The constant stream of sensory information provided by the environment must be filtered by the brain to reduce redundancy and prioritize behaviourally relevant signals. In sensory cortex this is first achieved by adjusting the gain of neural responses over time-scales of seconds through adaptative mechanisms reflecting the recent history of the input^1–7^. Information accumulated over days can then induce more persistent changes in cortical function that shape behaviour. Two simple examples of these slower adjustments are habituation to a repeated stimulus and associative learning contingent on reward or punishment^8–12^. Changes in cortical processing across these different time-scales have been observed for several sensory modalities, including olfaction^13^, audition^14^ and vision^8,9,11^ but they have almost always been studied in isolation despite occurring simultaneously. As a result, it is unclear how the circuit mechanisms underlying simple forms of learning interact with fast and reversible adaptation^7^.

In layer 2/3 of primary visual cortex (V1) opposing forms of short-term adaptation have been identified: while some pyramidal cells (PCs) respond strongly at stimulus onset and then depress, others begin to respond weakly but then gradually sensitize over seconds^6^. Optogenetic manipulations and modelling demonstrate that this heterogeneity arises from variations in the *balance* of direct inhibition that PCs receive from PVs and SSTs; stronger PV input promotes depression while SST inputs promote sensitization^6,15^. On time-scales of days, repeated exposure to a stimulus causes habituation characterized by reduced behavioural and neural responses^8,10–12^, while pairing of a stimulus with reward or punishment often enhance responses^8,9,16,17^. How do these slow changes in excitatory representation interact with the dynamics of interneurons that govern fast adaptation?

To monitor changes in circuit dynamics during simple forms of learning we used calcium imaging to record responses in PCs and the three major classes of interneurons (VIP, SST and PV) while using the *same stimulus* to probe fast adaptation, habituation and associative learning. We then integrated the data in a rate-based circuit model that provided predictions of changes in connection strengths that could be tested using optogenetics^15^. The results identify dissociable circuit motifs for the two forms of learning. Habituation *reduces* disinhibition through the VIP→SST→PC pathway both by weakening feedback activation of VIPs and by reducing the strength of VIP→SST synaptic connections. Reward association *increases* disinhibition through the SST→PV→PC pathway by increasing the strength of SST→PV connections, while simultaneously reducing SST→PC inhibition. The decrease in the PV:SST input ratio to PCs that accompanied both forms of learning enhanced the sensitizing form of fast adaptation, consistent with the “balance” model previously shown to account for the variation in adaptive properties across the PC population^6^. These results identify changes in cortical circuits that link fast adaptation to simple forms of learning.

## Results

### Simple forms of learning shift PCs towards the sensitizing form of adaptation

To investigate interactions between adaptive dynamics and learning we recorded activity from layer 2/3 PCs using two-photon calcium imaging (Fig. 1A) and inferred spike rates using MLSpike^18^. Our standard stimulus was a drifting grating of high contrast (20°, 1 Hz, 100% contrast) presented for 10 s while the mouse ran on a treadmill (Fig. 1B)^6^. Animals were exposed to the stimulus 50-200 times per session across six sessions at intervals of 2-3 days and divided into two groups: a habituation group and a reward-association group which were also given water 0.25 s following each stimulus presentation (Fig. 1C).

**Figure 1.**
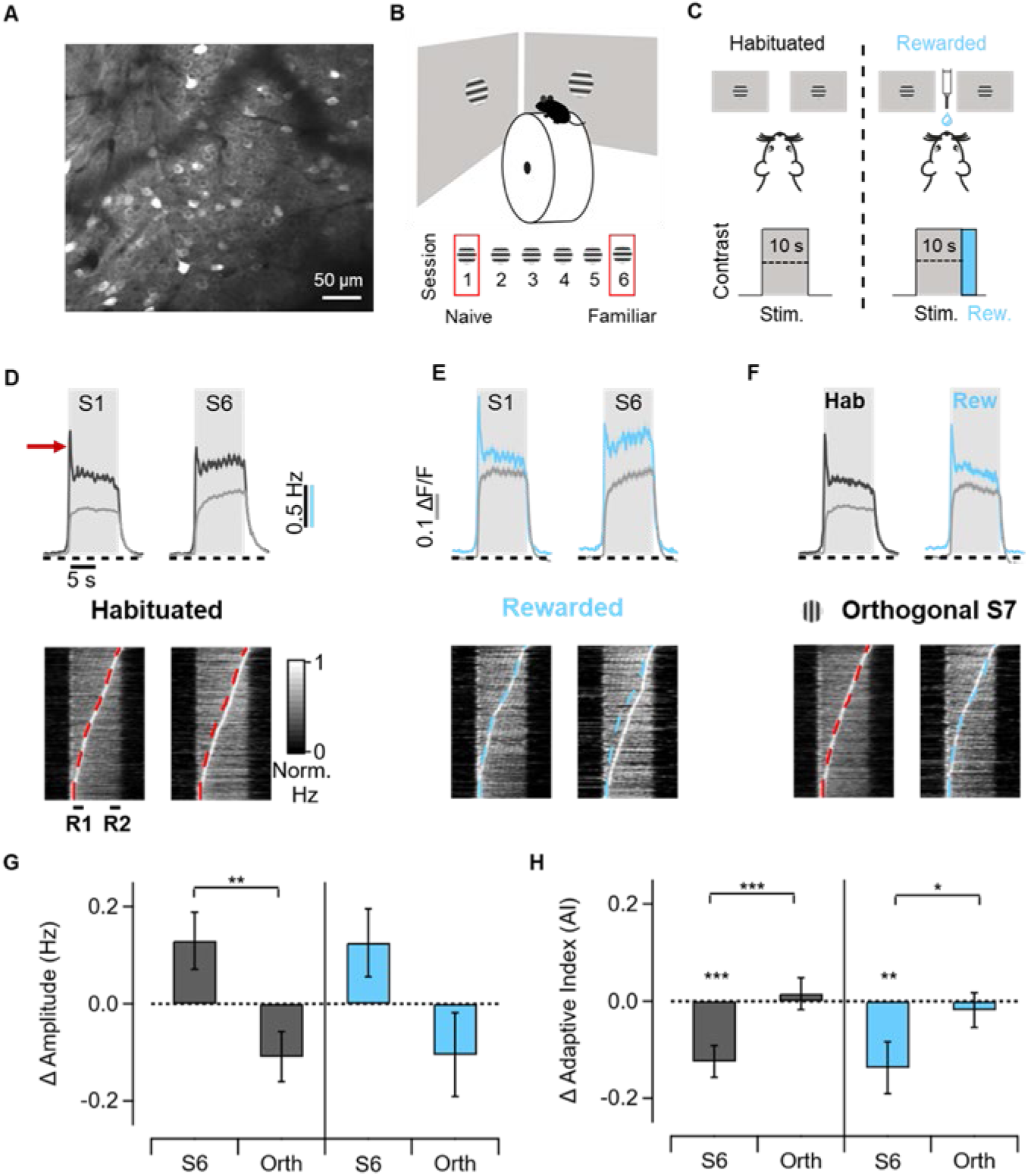
Stimulus familiarity alters PC adaptation. **A**. Average projection of an example field-of-view (FOV) in which PCs express GCaMP6f. **B**. Experimental set up. Mice head-fixed on a treadmill and free to run were presented with upwards drifting gratings over six sessions. **C**. Mice were divided in two groups to form different associations with the stimulus. The habituated group (left) was simply exposed to the stimulus while the rewarded group (right) was given water ∼0.25 s after each stimulus presentation. **D.** Top: Average PC responses in the habituated group to the stimulus in session 1 (S1) and session 6 (S6), shown as calcium signals (grey) and inferred spike rates (black). Averages include only PCs generating a significant positive response. Grey boxes indicate the 10 s stimulus presentation period, and the red arrow marks an initial depressing peak whose dynamics were not investigated further. Bottom: Raster plots of normalised PC responses sorted by time of peak activity. The dashed line in S6 retraces the trajectory of peak times in S1. R1 and R2 mark the amplitude window used to calculate adaptive indexes. (S1 1,444 Cells, S6 950 Cells from 14 mice) **E.** As D, but for the rewarded group of mice. (S1 525 Cells, S6 413 Cells from 6 mice) **F**. As D, but for mice presented with an orthogonal stimulus in session 7 (S7) (Hab. 1303 Cells, Rew. 524 Cells) **G**. Bar plot showing the change in response amplitude in S6 and S7 (orthogonal), both compared to S1. (Linear Mixed Models, LMM. Habituation 55 FOVs from 14 mice. Rewarded 30 FOVs from 6 mice. Interaction Hab S1- Rew S1, p = 0.44). Asterisks above bars indicate significant differences relative to S1, asterisks above connecting lines indicate differences between S6 and S7. **H.** As G, but for change in adaptive index. In both conditions, the AI significantly decreased over 6 sessions. *p < 0.05, **p < 0.01 and ***p < 0.01.

Two simple ways in which neural representations of a stimulus can be adjusted are changes in the number of neurons that respond and the amplitude of their responses^8,9,19^. The most obvious effect of habituation was a 33 ± 4% decrease in the number of responsive PCs, defined as those in which activity was significantly positively correlated with the stimulus (Table 1; see Methods). Of the PCs that did generate a significant response, the average amplitude over the 10 s stimulus was little affected by habituation (habituation ΔS1-S6 0.13 Hz, p = 0.2; Reward ΔS1-S6 0.13 Hz, p = 0.7; Fig. 1D,G). Notably, when the stimulus was paired with a water reward the density of PCs excited by the stimulus halved in session 1 compared to the habituation group and then stayed stable up to session 6 (Table 1). These PCs also generated larger responses, but again this was already evident from session 1 (Fig. 1E and G). These immediate effects of pairing the stimulus with water reward indicate a change in state which was also observed when the reward was dissociated in time from the stimulus (Fig. 1D-E; Supplementary Fig. 1A). The *specific* effect of associating the reward with the stimulus was to maintain a stable number of responsive PCs throughout the repeated sessions (Table 1, Supplementary Fig. 2A-B).

**Table 1.**
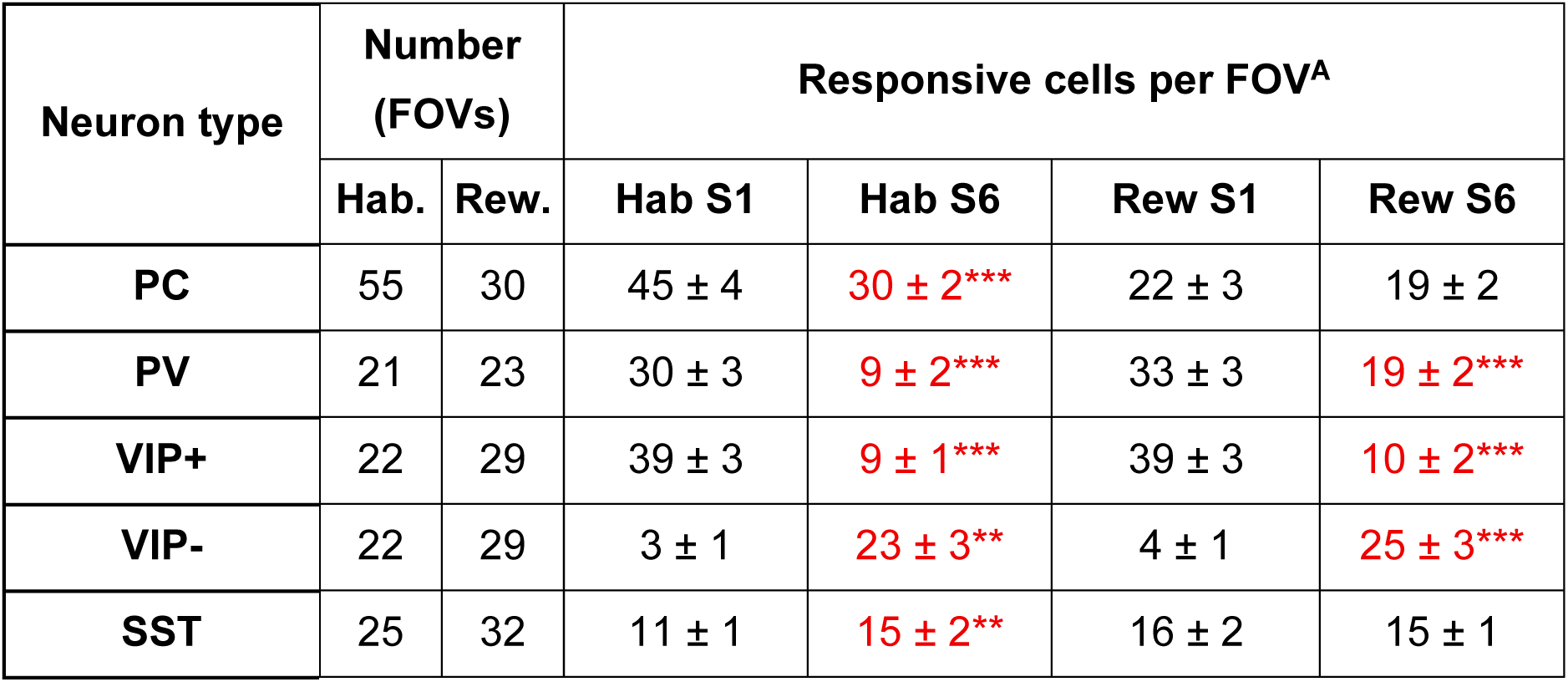
Reward association differentially regulates the number of responsive neurons. **A.** Number of stimulus-responsive neurons in Session 1 and Session 6 for each cell type per field of view (FOV). Mean ± SEM with significant changes highlighted in red (linear mixed-effects model (LMM), *p < 0.05, **p < 0.01 and ***p < 0.001). PCs decreased 33±4% with habituation while SST increased 36±6%, conversely none of them changed during reward association showing differential changes between both conditions (Session x Condition interaction, p < 0.05). PVs dramatically decreased with habituation (70±17% decrease) and to a lesser extent with reward association (42±6%). Lastly, while VIP+ experienced a drastic reduction (77±10 % Hab, 74±16% Rew) VIP-increased dramatically in both conditions (667±239 % Hab, 525±146% Rew) (Number of mice: PC Hab 14, PC Rew 6, PV Hab 3, PV Rew 4, VIP Hab 3, VIP Rew 4, SST Hab 4, SST Rew 5).

| Neuron type | Number (FOVs) |  | Responsive cells per FOV<sup>A</sup> |  |  |  |
| --- | --- | --- | --- | --- | --- | --- |
|  | Hab. | Rew. | Hab S1 | Hab S6 | Rew S1 | Rew S6 |
| PC | 55 | 30 | 45 ± 4 | 30 ± 2*** | 22 ± 3 | 19 ± 2 |
| PV | 21 | 23 | 30 ± 3 | 9 ± 2*** | 33 ± 3 | 19 ± 2*** |
| VIP+ | 22 | 29 | 39 ± 3 | 9 ± 1*** | 39 ± 3 | 10 ± 2*** |
| VIP- | 22 | 29 | 3 ± 1 | 23 ± 3** | 4 ± 1 | 25 ± 3*** |
| SST | 25 | 32 | 11 ± 1 | 15 ± 2** | 16 ± 2 | 15 ± 1 |

An important difference between the two groups of mice was that the reward group had limited water between recording sessions. The resulting weight loss can affect neural responses^20^ raising the possibility that changes in the number of responsive PCs in the rewarded and habituated groups are a side-effect of water deprivation rather than formation of a reward-stimulus association. Evidence against this idea was provided by measurements in a control group of water-deprived mice in which the reward was provided uncoupled in time from the stimulus: these mice still exhibited a reduction in the number of responsive PCs across sessions (Supplementary Figure 1A-C). This reduction was correlated with a failure to associate the stimulus with the reward, as assessed by the absence of two behavioural indicators: a sudden acceleration in locomotion at stimulus onset (Supplementary Figure 1D-E) and a progressive dilation of the pupil while the stimulus was maintained (Supplementary Figure 1F-G).

To investigate how these changes in PC responsivity might relate to adaptation during the 10 s stimulus we measured an adaptive index, AI = (R1 - R2) / (R1 + R2), where R1 and R2 are the average response over 2 s time windows at the start and end of the stimulus respectively (Fig. 1D). R1 was measured with a delay of 1 s to avoid the initial very fast depressing component of the response reflecting adaptation in the feedforward input (arrowed in Fig. 1D). Positive AI indicates depression and negative AI indicates sensitization. Across sessions, the average response of the PC population gradually shifted from mild depression toward sensitization in both the familiarity-only and reward-associated groups (Fig. 1D,E & H; Supplementary Fig. 2C,D). All these changes were stimulus specific: presentation of an orthogonal grating restored response dynamics resembling session 1 (Fig. 1F-H, Supplementary table 1). The adaptive dynamics within the PC population therefore shift towards sensitization when a stimulus becomes familiar, whether or not the stimulus is made salient by an associated reward.

### Reorganization of inhibition during simple forms of learning

What are the circuits that shift PCs towards the sensitizing form of adaptation during these simple forms of learning? Inhibitory neurons in V1 have been shown to have more pronounced adaptive properties than PCs, with PV and VIPs being predominantly sensitizing while SSTs are mostly depressing^6,15^ (Fig. 2). As a result, a decrease in the ratio of PV to SST inputs favours sensitizing adaptation in PCs^21^. We therefore measured the effects of the habituation and reward protocols on responses in the three major interneuron populations – PV, SST and VIP.

**Figure 2.**
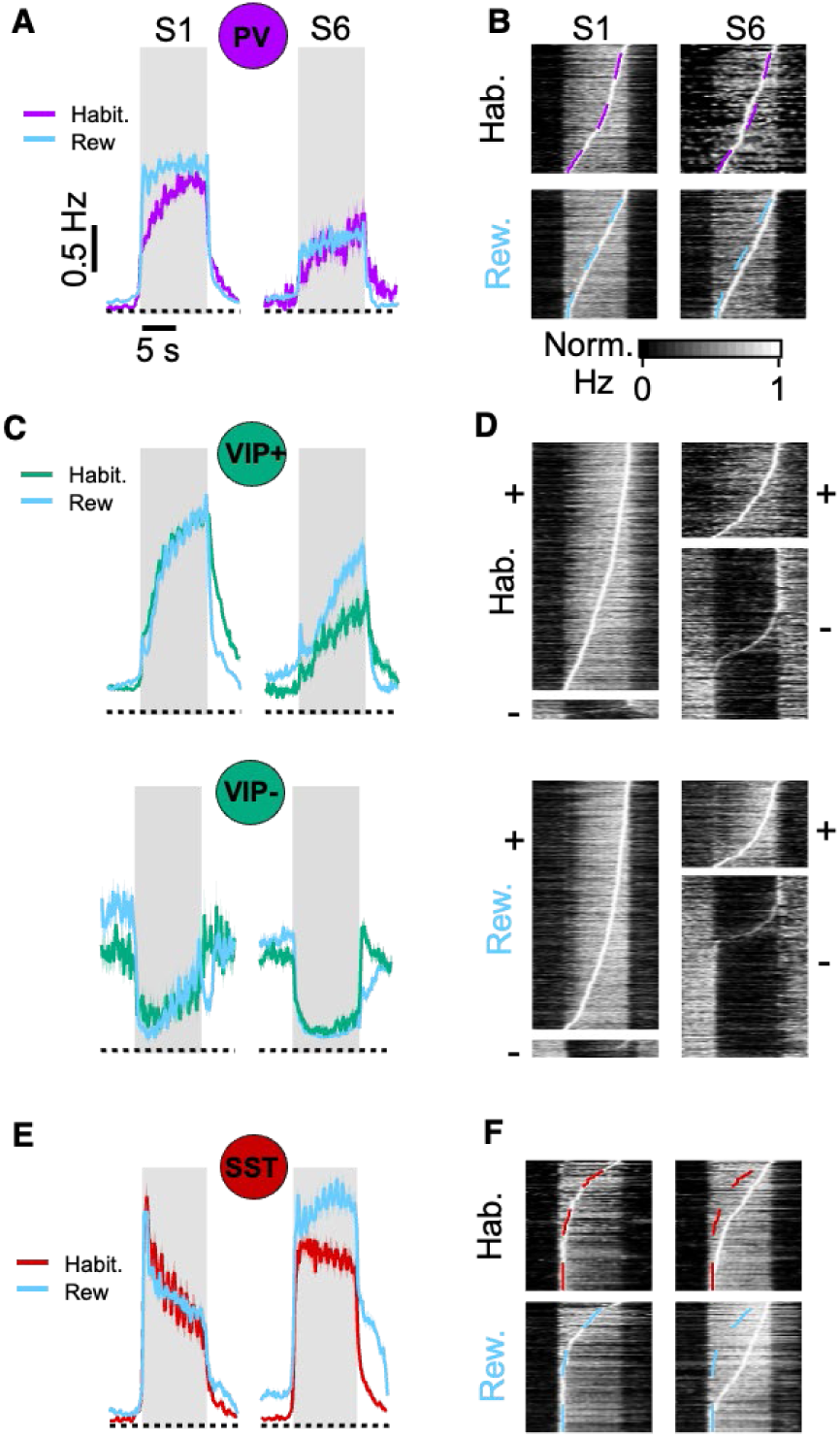
Changes in interneuron activity during two simple forms of learning. **A.** Average PV interneuron responses in S1 and S6 of habituated (purple) and rewarded (blue) animals. These are only averaged from PVs generating a significant positive response. Responses were suppressed in both habituated and rewarded groups of mice (Habituation 46 ± 7% reduction. Reward 53 ± 6% reduction. LMM, both p < 0.001. See Supplementary Figure 3). **B.** Raster plots of normalised responses sorted by time of peak activity. The dashed lines retrace the peak times of S1 in either condition, showing that adaptive properties of the population remained similar. The distribution of peak times of individual neurons stays constant irrespective of reward. (Habituation: S1 427 Cells, S6 91 Cells from 3 mice. Reward: S1 572 cells, S6 261 cells from 4 mice) **C**. As A, but for VIP interneurons which were divided in two populations: those responding positively to the stimulus (VIP+ green, top) and those suppressed by the stimulus (VIP- green, bottom). Notice the decrease in amplitude in S6. (VIP+. Habituation 55 ± 5% reduction. Reward 33 ± 9% reduction. LMM, both p < 0.001. See Supplementary Figure 3). **D.** As B, but for VIP+ and VIP- cells. Note the large increase in the number of VIP- cells at the cost of the number of VIP+ cells (quantified in Table 1) (Habituation: S1 757 VIP+, 59 VIP- Cells; S6 VIP+ 178, VIP- 321 Cells from 3 mice. Reward: S1 VIP+ 950, VIP- 67 cells, S6 VIP+ 207, VIP- 436 cells from 4 mice) **E, F.** As A, but for SST interneurons (Habituation: S1 219 Cells; S6 291 Cells from 4 mice. Reward: S1 291 cells, S6 312 cells from 5 mice). Note the increased amplitude of responses in the habituated and rewarded groups (Habituation 80 ± 19% increase. Reward 86 ± 16% increase. LMM, both p < 0.001). and the shift away from depressing adaptive dynamics towards sensitization (ΔAI. Habituation -0.23 ± 0.06, LMM p < 0.01. Reward -0.19 ± 0.05, LMM p < 0.001. See Supplementary Figure 3).

PV interneurons exert powerful perisomatic inhibition onto PCs to control excitatory gain in V1^22,23^. Despite this general role, the habituation-induced decline in PC responsivity (Table 1) was accompanied by a large reduction in PV interneuron recruitment (70 ± 17% in habituated mice; 42 ± 6% in rewarded mice; Table 1; Fig. 2A-B). These results align with previous reports of reduced PV activity upon exposure to familiar stimuli in V1^24^ and immediately suggest that the other source of direct inhibitory inputs to PCs, SST interneurons, counterbalance the reduction in PV activity. This was confirmed by directly observing activity in SSTs: in the habituation group of mice there was a 36 ± 6% increase in the number of SST interneurons that were significantly responsive to the stimulus, but this effect was completely abolished by reward association (Table 1). In both habituated and rewarded groups SSTs also underwent a shift from depressing to sensitizing forms of fast adaptation (Fig. 2E-F; habituation AI shift from 0.34 ± 0.05 to 0.11 ± 0.04; reward association AI shifts from 0.26 ± 0.04 to 0.07 ± 0.03), paralleling the familiarity-driven changes observed in PCs (Fig. 1). Changes in the dynamics and responsivity of PV and SST interneurons can therefore be expected to lead to a decrease in the PV:SST ratio of PCs, consistent with the familiarity-driven shift toward sensitizing adaptation that was observed.

VIP interneurons are a major target of long-range signals arriving in V1 from the basal forebrain as well as feedback from higher visual areas ^25,26^. In naïve mice, they fell into two distinct functional subpopulations: the large majority (93 ± 2%) were relatively quiescent at rest and sensitized during high-contrast stimulus presentation (VIP+), whereas a small minority (7 ± 2%) exhibited higher spontaneous activity that was *suppressed* by the stimulus (VIP-; Table 1; Fig. 2C-D) ^27,28^. The VIP- subpopulation expanded dramatically in habituated mice, up to 71 ± 5% of all responsive VIPs, and a similar shift was observed in rewarded mice. This large increase in the density of VIPs suppressed by the stimulus was, unsurprisingly, simultaneous with a decrease in the density of VIPs excited by the stimulus. Although stimulus-suppressed VIP responses have been described previously^27,28^, their potential role in experience-dependent plasticity has not been clear. These results demonstrate that learning to recognize a familiar stimulus is associated with a profound reorganization of VIP circuitry, irrespective of the salience of that stimulus.

These results demonstrate that as a stimulus becomes familiar, adjustments in signal flow through layer 2/3 involve all major cell types to varying degrees. One of the most powerful changes is within the VIP population, where the ratio of neurons excited by the stimulus compared to those inhibited changed from 13 in session 1 to 0.4 in session 6. Even among VIP+ group, the average response fell by a factor of ∼0.4 (Fig. 2C, Supplementary Fig. 3A2). VIPs are a major target of external inputs arriving to layer 2/3 from other cortical areas, such as frontal or higher-order sensory cortices ^25,29,30^, which adjust PC responses through the VIP→SST→PC disinhibitory pathway. Similar changes in VIP responses occurred in the rewarded group of mice, yet the density of responsive PCs did not change from session 1 to session 6 (Table 1). Below we show how reward association compensates for decreased VIP activity through a circuit readjustment that recruits a second disinhibitory pathway, SST→PV→PC, thereby preventing the loss of responding PCs across repeated stimulation sessions.

### Distinct disinhibitory pathways for familiarity and reward

A model is required to quantitatively relate the activity of different types of neurons because these are interconnected through multiple pathways with different connection weights (Fig. 3A). The task of comparing naïve, habituated and reward-associated groups of animals is made even more complicated by the fact that the strength of individual synaptic connections is not fixed, but subject to modulation^15,31^, for instance through presynaptic receptors or by post-synaptic modifications^32–34^. Such changes in the strength of individual synaptic connections have been very hard to quantify in behaving mice because recording from connected neuron pairs requires anaesthetics that disrupt inhibition^35–37^. We used a circuit model that provides predictions of the changes in connection strengths occurring during habituation and reward association^15^ (Figs. 3-4) and then tested these predictions by optogenetic manipulation of the different interneuron classes (Figs. 5-6).

**Figure 3.**
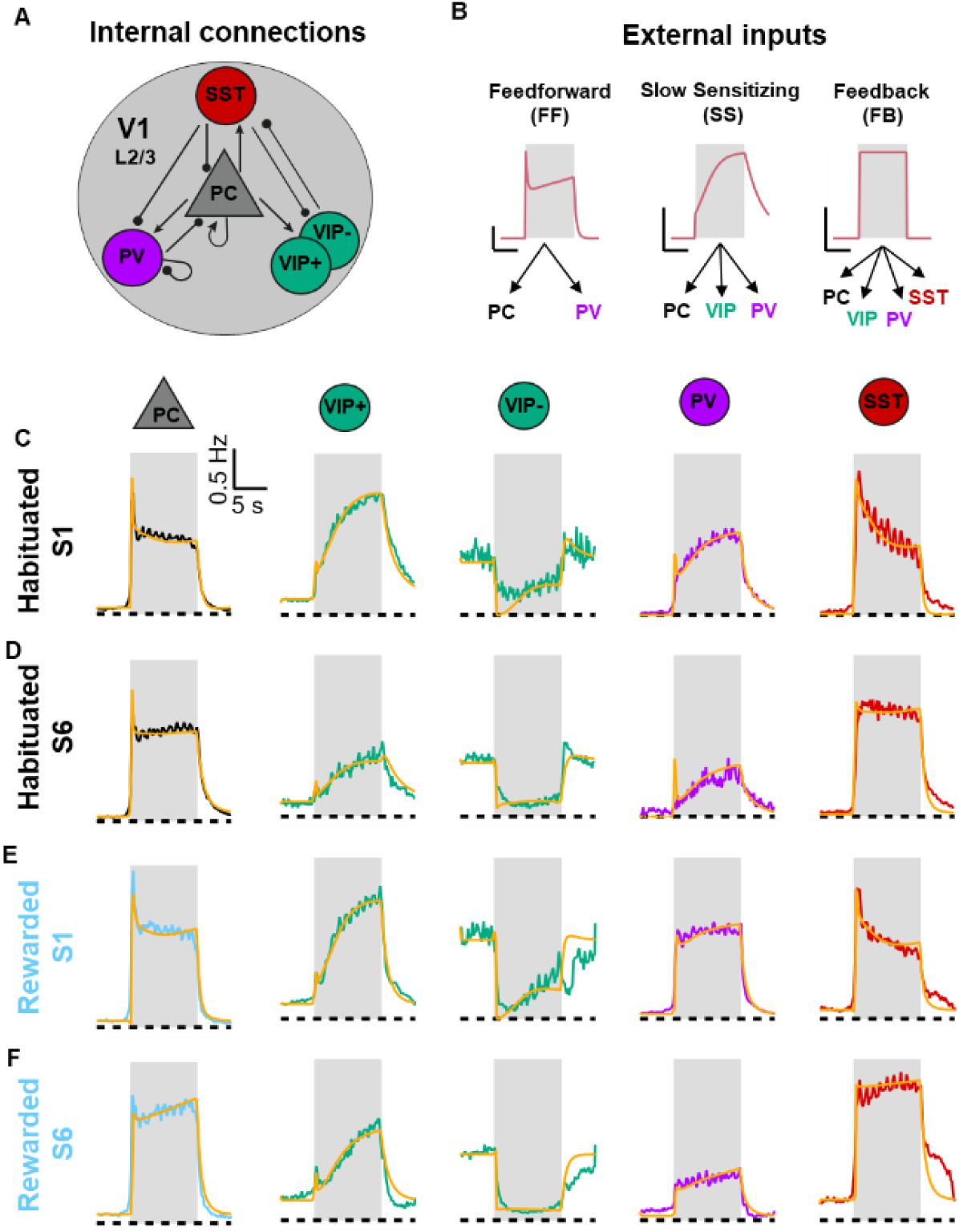
A data-driven circuit model reproduces patterns of circuit activity in naïve, habituated and rewarded mice. **A.** Schematic of the model showing the major excitatory (arrow tip) and inhibitory (round tip) local connections in layer 2/3 of V1. **B.** Three external inputs drove activity: feedforward excitation (FF); feedback input (FB) and a slow sensitizing input (SS). Their connections to V1 neurons are shown. (Vertical bar: 0.5 Hz, horizontal bar: 5 s) **C, D**. Average firing rates of PCs and four types of interneurons (VIP+ Green, VIP- Green, PV purple, and SST red) during stimulus presentation and their corresponding fits calculated by the model (yellow) for S1 and S6 of habituated mice. **E, F**. As C and D but for mice in the rewarded group.

**Figure 4.**
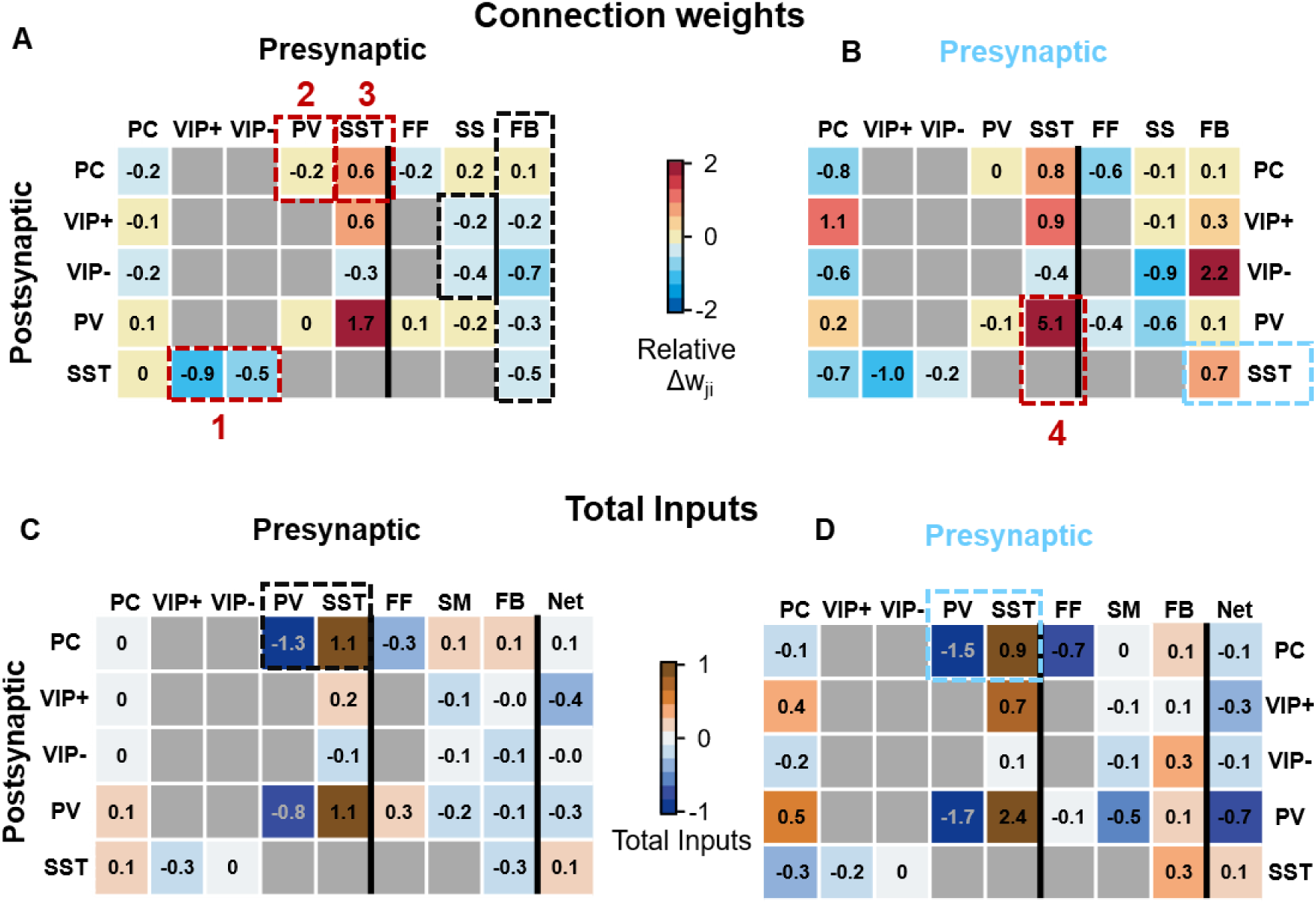
Distinct circuit mechanisms underly habituation and reward association. **A.** The relative change in connection weights caused by habituation from session 1 to session 6. Note the decrease in VIP->SST connections, the increase in SST connections to most targets, and, in the external inputs, the decrease in FF, SS and FB. **B.** same as in A for rewarded mice. Note the increase in SST to PV connection and FB that were different to the habituated mice. **C-D**. The change in the total synaptic input activated by the stimulus, calculated from the average activity and the change in connection weight in A-B (See Methods). The last column is the net change taking into account the polarity of the synapse. Notice the dramatic decrease in PV inputs and increase in SST inputs in both habituated and rewarded models (black and blue boxes).

**Figure 5.**
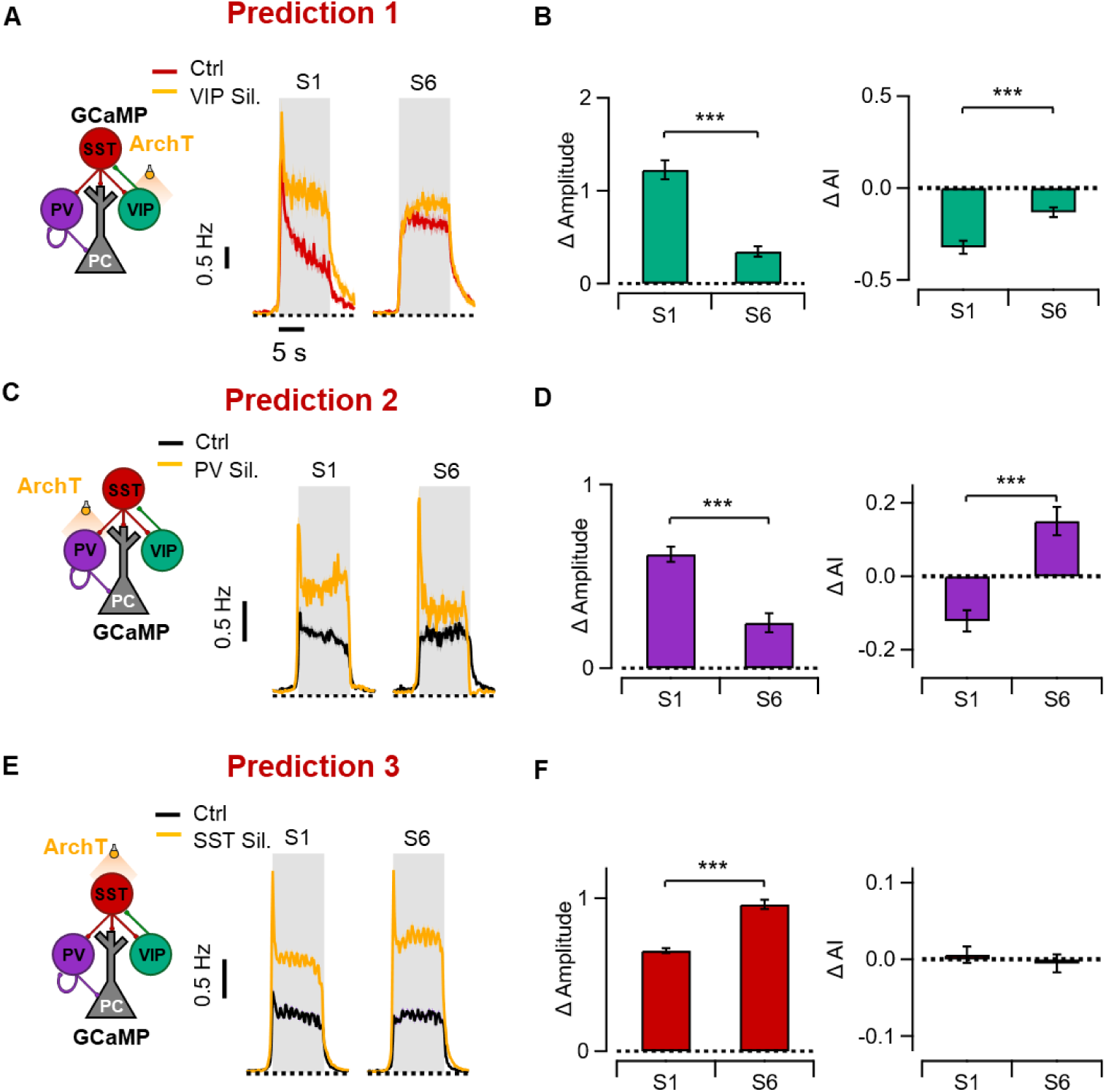
Optogenetic tests of changes in circuit connectivity during habituation. **A.** Left: Schematic of optogenetic manipulation testing prediction 1 (reduction in VIP->SST connectivity). During stimulus presentation VIPs were inhibited with ArchT while we simultaneously imaged SSTs expressing GCaMP6f. Right: average SST responses of habituated animals to the stimulus in S1 (84 cells, 4 mice) and S6 (104 cells) in control trials (red) and trials paired with VIP inhibition (yellow). **B.** Bar graph showing decrease in average response (left) and increase in AI (right) of SSTs when suppressing VIPs (experiment in A). Both effects were reduced by habituation. **C.** Left: Schematic of optogenetic manipulation testing prediction 2 (reduction in PV->PC connectivity). During stimulus presentation PVs were inhibited with ArchT while we monitored GCaMP responses in PCs. Right: average PC responses of habituated animals to the stimulus in S1 (446 cells, 4 mice) and S6 (258 cells) in control trials (black) and trials paired with PV inhibition (yellow). PV suppression led to an increase in PC response gain that was stronger in S1 compared to S6, indicating a reduced PV influence over PCs during habituation. **D.** The change in average response (left) and AI (right) of PCs during experiment in C. Following optogenetic stimulation the AI shifted towards sensitization in S1 while becoming more depressing after habituation, and the decrease in amplitude was stronger in S1. **E.** Left: Schematic of optogenetic manipulation testing prediction 3 (enhanced SST- >PC connectivity). During stimulus presentation SSTs were inhibited with ArchT while we monitored GCaMP responses in PCs. Right: average PC responses of habituated animals to the stimulus in S1 (1243 cells, 5 mice) and S6 (890 cells) of control trials (black) and trials paired with SST inhibition (yellow). **F.** The change in average response (left) and AI (right) of PCs during experiment in **E**. SST suppression led to an increase in PC response gain that was significantly stronger in S6 compared to S1, indicating strengthen SST influence over PCs during habituation. AI remained unchanged. Linear mixed model (LMM) ***p < 0.01.

**Figure 6.**
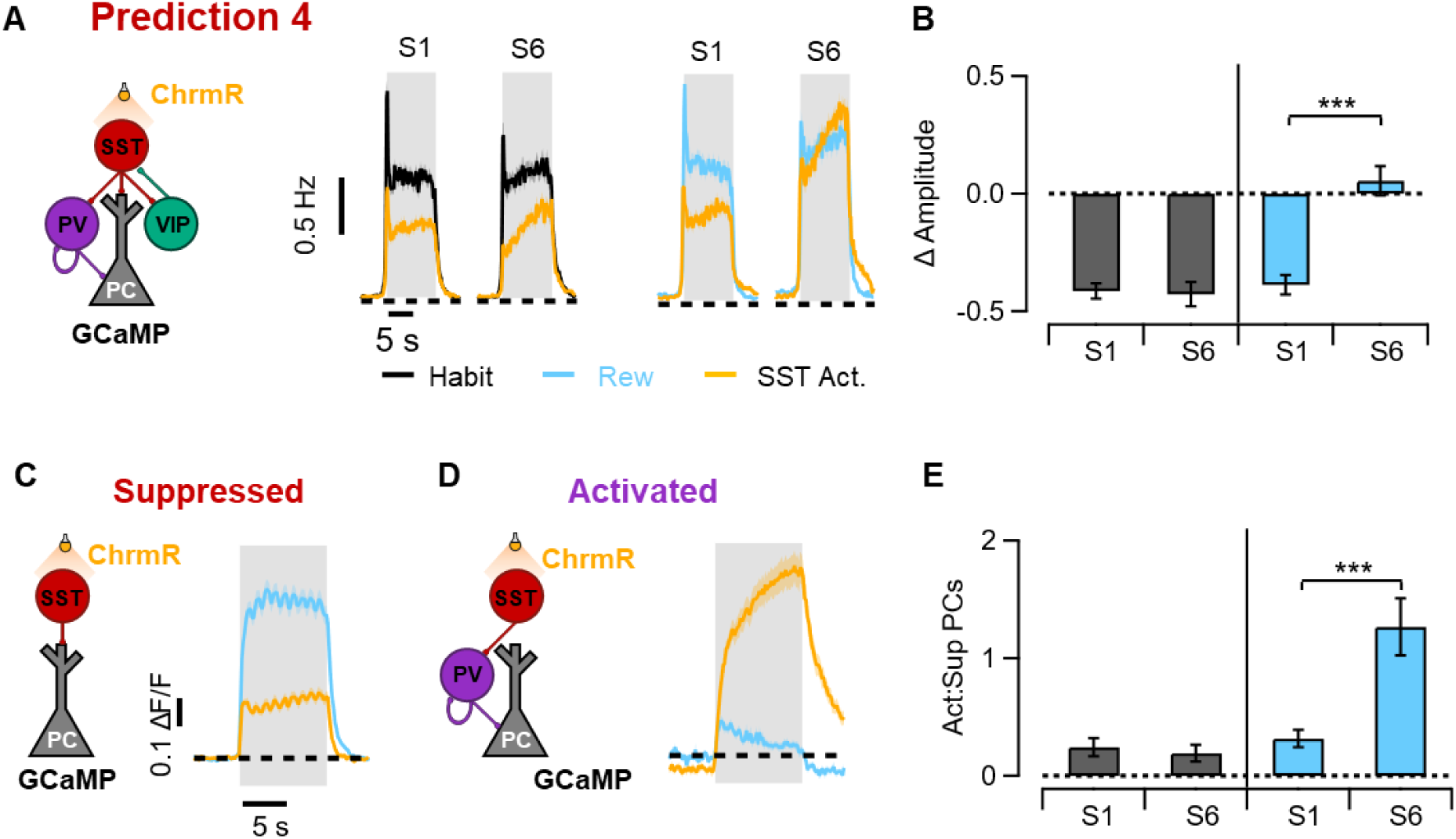
Reward association alters SST inhibition. **A**. Left: Schematic of optogenetic manipulation testing prediction 4 (Increase in SST->PV input). During stimulus presentation SSTs were activated with ChrimsonR while PCs expressing GCaMP6f were imaged. Middle: Average PC responses of habituated animals to the stimulus in S1 and S6 of control trials (black) and trials paired with SST activation (yellow). Right: PC responses of rewarded animals (blue: control, yellow: paired with SST activation). **B.** Bar graph showing the change in amplitude of PCs when trials were paired with LED illumination to activate SSTs during S1 and S6. Whereas the change in amplitude remained stable across sessions during habituation it decreased significantly in rewarded animals, indicating weaker SST-dependent inhibition. (LMM. Significant at p <0.001) **C.** Average response of the PC sub-population that was suppressed under SST activation and schematic showing the dominant pathway - direct inhibition from SSTs. This average is only from the rewarded group. **D.** Average response of the PC sub-population that was activated under SST activation and schematic showing the dominant pathway - SST→PV→PC disinhibition. **E.** Bar chart showing the relative number of PCs that were activated or suppressed by inhibiting SSTs. In naive and habituated mice suppressed PC outnumbered activated by a factor of ∼4, indicating dominance of direct SST→PC inhibition. But after reward association activated PCs outnumbered suppressed by a factor of ∼1.3, indicating that the SST→PV→PC disinhibitory pathway had become dominant (LMM. Significant at p <0.001).

The model considers all the major internal connections and external inputs to layer 2/3 and has previously been shown to account for rapid changes in circuit function associated with locomotion^15^ (see Methods). Briefly, it consists of a set of differential equations where the connection weight between presynaptic population j and postsynaptic population i is represented by w_ji_ (Fig. 3A), which is simply the product of the total number of synaptic connections, *C_ji_*, and the average strength of an individual synapse between neuron type j and i, *s_ji_*:

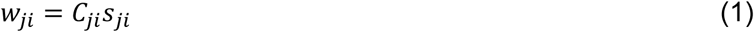

External inputs are a feedforward signal from layer IV (FF), feedback from higher order cortices (FB), and a slow sensitizing signal (SS) that strongly targets VIP and PV interneurons (Fig 3B). We made one significant change from the previous form of the model by considering VIP interneurons to be of two subtypes, those positively correlated with the stimulus (VIP+) and those negatively correlated (VIP-), based on the clear functional distinction between these two groups shown in Fig. 2C-D. Fitting the model produced sets of solutions that described the response dynamics of PCs and interneurons, with the average connection weights (*w_ji_*) being representative of the distribution of solutions for each connection (Fig. 3C-F, see Methods). Values of *_ji_* in naïve mice (session 1) are shown in Supplementary Figure 4 and the relative changes between sessions 6 and session 1 (equation 2) are shown in Fig. 4A for the habituated group of mice and in Fig. 4B for the rewarded mice, calculated as

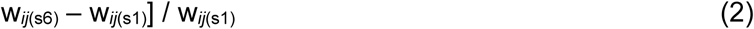

Habituation altered the matrix of estimated connection strengths in three general patterns (dashed boxes in Fig. 4A), all impacting the VIP→SST→PC disinhibitory pathway: i) reduced external feedback (FB) to VIP and SST interneurons, and reduced SS inputs to VIPs (Δw_FB→VIP+ and VIP- ≈ -20% and -70%; Δw_FB→SST ≈ -50%; Δw_SS→VIP+ and VIP- ≈ -20% and -40%); ii) stronger SST outputs to multiple targets and particularly PVs (Fig. 3C and D; Fig. 4A, Δw_SST→PC ≈ +60%; Δw_SST→PV ≈ +170%; Δw_SST→VIP+ ≈ +60%), and iii) weakened VIP→SST inhibitory connections (Δw_VIP+→SST ≈ -90%; Δw_VIP-→SST ≈ -50%). Reduced feedback is consistent with a range of previous experiments showing that top-down drive to primary sensory cortex falls as the stimulus becomes familiar and predictable so we did not test these predictions further^8^. Habituation-dependent changes in connectivity within layer 2/3 have been harder to assess and the model predicted distinct changes in the inputs and outputs to SSTs and PVs, as highlighted by the three red boxes in Fig. 4A. Below, these three predictions are tested using optogenetics and found to hold (Fig. 5 below), indicating that the suppression of familiar information that is not behaviourally relevant occurs through two coordinated adjustments; reducing excitatory feedback from higher visual areas and reduced VIP-mediated disinhibition of PCs.

When the stimulus was associated with reward, the model predicted that the preservation of PC responsivity occurred through a distinct circuit motif - the SST→PV→PC disinhibitory pathway^38,39^. This mechanism of gain control became dominant over direct SST→PC inhibition through two key changes (Fig. 4B): a marked increase in the strength of feedback signals to SSTs (Δw_FB→SST ≈ +70%; blue box in Fig. 4B), and a much stronger SST connection to PVs (Fig. 4D; Δw_SST→PV, ≈ +500%; red box 4). The first change is again consistent with a range of experiments showing that top-down drive to SSTs increases as a stimulus becomes associated with a reward^8^, and was not tested further. But strengthening of SST→PV connections has not been identified previously and was therefore tested using optogenetics when it was found to hold (Fig. 6 below). In parallel, reward association limited SST→PC inhibition by preventing the habituation-induced increase in SST recruitment(Fig. 2E-F and Table 1). In other words, reward shifted the relative importance of the two pathways controlling PC activity in response to external inputs through SSTs, away from the VIP→SST→PC pathway and towards the SST→PV→PC pathway.

### Optogenetic tests of changes in circuit connectivity during habituation

The model predicts that habituation reduced VIP-mediated disinhibition of PCs through changes in the strength of at least three connections within the circuit, numbered in Fig. 4A. These predictions were tested using optogenetics.

*Connection 1: VIP→SST.* VIP interneurons were inhibited using ArchT while monitoring GCaMP activity in their primary target, SSTs. Stimulus trials in which the LED was switched on were interleaved with control trials and the average responses are shown in Fig. 5A for sessions 1 and 6. In naïve mice, VIP silencing markedly increased SST gain and shifted the population towards sensitizing adaptation, demonstrating a strong connection. Both these effects were abolished by session 6, indicating a significant weakening of this connection by habituation (Fig. 5B; p < 0.001), as predicted by the model (box 1 in Fig. 4A).

*Connection 2: PV→PC.* Inhibiting PVs using ArchT caused strong sensitization and gain increases in PCs in session 1, but these were almost completely abolished by session 6 (Fig. 5C-D; p < 0.001), consistent with the observed PV disengagement during habituation (Fig. 2A; Table 1) and the fit of the model (box 2 in Fig. 4A).

*Connection 3: SST→PC.* SST inhibition increased PC gain in naïve mice, and this was *potentiated* after habituation, indicating a strengthening of this connection (Fig. 5E-F; p < 0.001). This result is in line with an increase in SST responsivity (Table 1) and was predicted by the model (box 3 in Fig. 4A).

This combination of experiments and modelling demonstrates that habituation causes a general switch in the source of direct inhibition to PCs, away from PVs towards SSTs. The model can also help us quantify this effect as the change in the *total* direct synaptic input that PCs receive from PVs and SSTs as they respond to the visual stimulus. The PV:SST input ratio fell by a factor of 1.2 in the habituated group of mice (black box in Fig. 4C) and a factor of 1.7 in the rewarded group (blue box in Fig. 4D). Using the “balance model”^6^, the shift towards the sensitizing form of adaptation during reward association can then be understood as reflecting this decrease in the PV:SST input ratio.

### Reward association strengthens the SST-PV-PC disinhibitory pathway

SST interneurons were the only neuronal population exhibiting specific and strong reward-dependent modulation in their response to the stimulus (Table 1; Fig. 2C). Similarly, they were the only type for which the model predicted a reward-selective change in the connection weights within layer 2/3 – a 3-fold increase in the strength of SST→PV connections compared to habituated mice (dashed red box 4 in Fig. 4B). To test this prediction, we used ChrimsonR to over-activate SST interneurons during stimulus presentations while imaging PC activity, again interleaving optogenetic and control trials (Fig. 6A). SST activation in naïve mice caused a net inhibition of ∼41%, indicating that direct SST→PC connections outweighed the effects of activating the SST→PV→PC disinhibitory circuit. This dominance of direct inhibition was maintained throughout sessions in the habituated group but almost abolished if the stimulus became associated with a reward, demonstrating a decrease in the inhibitory influence of SSTs (Fig. 6A and B; ∼5% increase; p < 0.001).

The results in Fig. 6A do not identify a single connection linked to reward association because SSTs influence the activity of PCs both directly and indirectly. To characterise this effect further, we separated all PC responses according to the consequences of the optogenetic manipulation. Fig. 6C shows average responses in PCs which were significantly *suppressed* by activating SSTs, reflecting dominance of the direct SST→PC connection, and Fig. 6D shows average responses in PCs that were *activated* by the optogenetic manipulation, reflecting dominance of the SST→PV→PC disinhibition. Notably, in the activated population of PCs the AI was strongly sensitizing upon SST activation (AI = -0.18 ± 0.03). In naïve and habituated mice, suppressed PCs outnumbered activated by a ratio of ∼4:1 (Fig. 6E), demonstrating the dominance of direct SST inhibition. Strikingly, after reward-association, activated PCs outnumbered suppressed by a ratio of 1.3:1. The model’s estimation of ∼3-fold strengthening of SST→PV synaptic connections with reward (prediction 4 Fig. 4B) therefore provides a precise mechanistic explanation for both the reward-specific increase in gain and the associated shift towards the sensitizing mode of fast adaptation.

## Discussion

The mechanisms that adjust sensory processing on different time-scales have typically been studied in isolation, leaving it unclear how far slower changes associated with learning reconfigure the inhibitory mechanisms that set the direction and magnitude of short-term adaptation^6,14,24,40^. Here we addressed this problem by probing fast response dynamics with an identical visual stimulus across days while independently manipulating stimulus relevance through reward. Three principal findings emerge. First, familiarity shifted PC population from weak depression toward sensitization, and this shift occurred whether or not the animal learnt to associate the stimulus with a reward (Fig. 1). Second, habituation reduced the fraction of responsive PCs while reward association preserved responsivity (Table 1). Third, the salience of the stimulus differentially adjusted two disinhibitory pathways that adjust PC gain in response to long-range input: while habituation *reduced* disinhibition through the VIP→SST→PC pathway, reward association *increased* disinhibition through the SST→PV→PC pathway. Using a combination of modelling and optogenetics we identify changes in connectivity within layer 2/3 circuitry that underly this plasticity, which are summarized in Fig. 7. Habituation weakened activation of VIPs while the stimulus was presented (Fig. 2–3) and reduced the strength of VIP→SST synaptic connections (Figs. 5–6). Associating the stimulus with a reward increased the strength of SST→PV connections and weakened SST→PC inputs (Figs. 4–6).

**Figure. 7.**
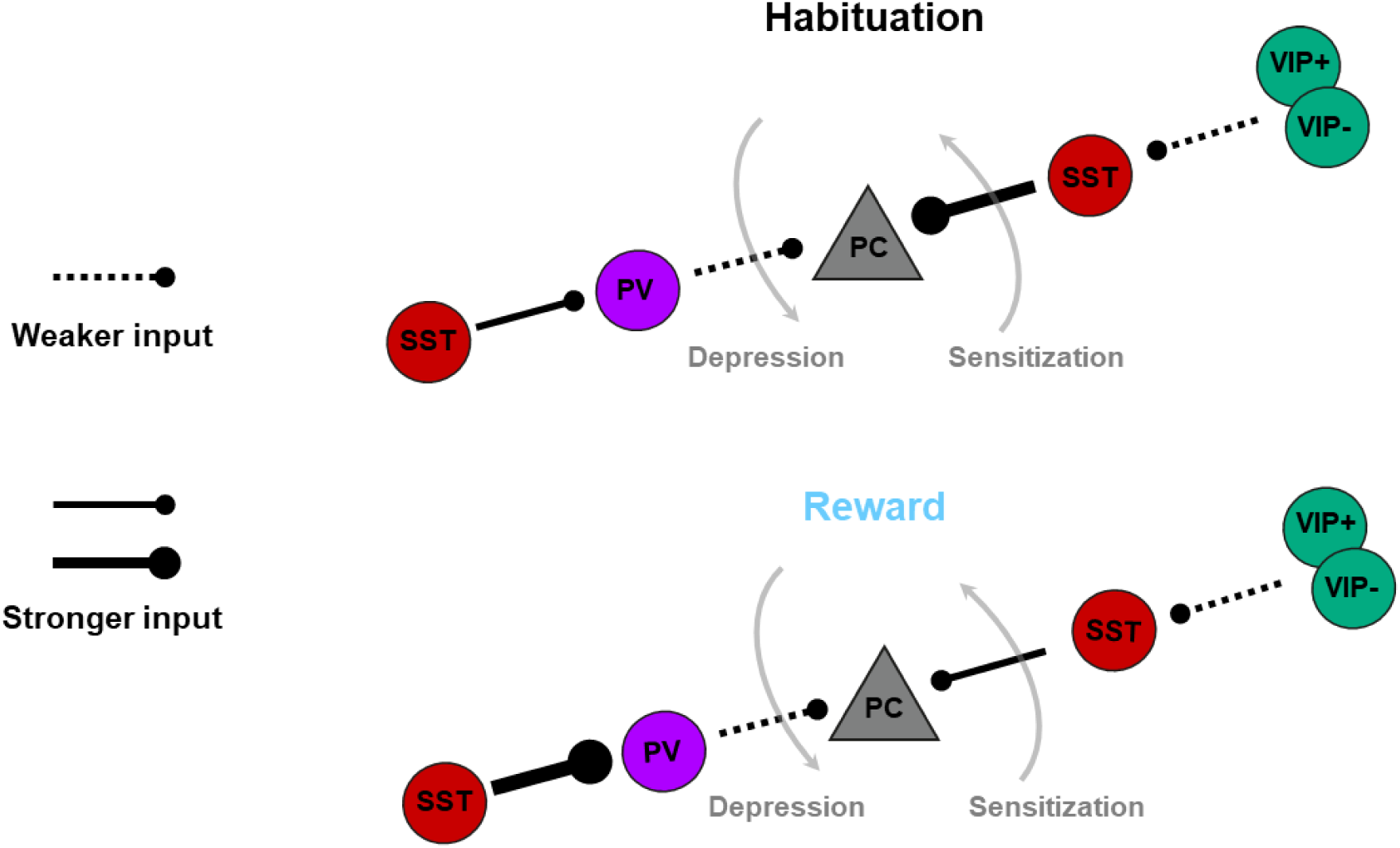
Summary of circuit reorganization driven by familiarity and reward association. Repeated visual experience engages two different inhibitory circuits based on distinct properties of the stimulus: familiarity and reward association. Familiarity, present in both habituated and rewarded mice, similarly reduces PV and VIP inhibition. In contrast, reward association prevents a larger increase in SST->PC direct inhibition while boosting SST->PV inhibition, shifting the balance from VIP->SST->PC to SST->PV->PC-dominated disinhibition.

A major determinant of the time-scale of visual adaptation in layer 2/3 is an external input that is slowly sensitizing and potentiated by changes in internal state associated with the onset of locomotion^15^. This input targets VIP and PV interneurons and is gated by cholinergic signals (possibly from secondary cortex LM^41^). Here we show that excitation of VIPs by these inputs is strongly reduced as a stimulus becomes familiar, whether or not it is also salient (Figs. 2 and 4). Optogenetic silencing of VIPs also shows that VIP→SST synapses are weakened by habituation (Fig. 5A). The combined effect of these changes was to increase SST activation and decrease PV responses (Fig. 2), driving the PC population to sensitizing adaptation.

The expansion of stimulus-suppressed VIP- neurons from ∼7% to ∼70% with habituation was striking. Previous work has described VIP- cells but this degree of plasticity within the VIP population has not been documented previously. Because VIP- cells are a small minority in naïve mice, the VIP population has been treated as homogenous in studies investigating fast state-dependent modulation^15,26,38^. Here we show that the large increase in the VIP- population correlates with an increase in the strength of the inhibitory SST→VIP connection (Fig. 6), highlighting these synapses as a key site of plasticity on longer time-scales.

What is the role of sensitization in signalling novelty or familiarity? Sensitizing neurons are better suited to detecting decreases in contrast or stimulus offsets, which may signal the end of a predictable event and cue environmental changes requiring attention. In contrast, depressing neurons, dominant for novel stimuli, provide rapid, transient responses ideal for detecting stimulus onsets or increases in contrast, facilitating reactions to unexpected inputs. These characteristics align in a general way with predictive coding frameworks, where novel stimuli represent prediction errors demanding fast behavioural responses, while familiar stimuli allow for more sustained monitoring of deviations from expectations. Our results extend earlier findings on SST upregulation for irrelevant stimuli^8,9,14^ by quantifying the underlying synaptic adjustments and linking them to adaptation.

Whereas simple habituation suppresses PC responses through increased SST inhibition (Table 1; Fig. 4 and 5), associating the stimulus with a reward prevented this not only by reducing SST→PC inputs but also by strengthening the SST→PV→PC disinhibitory pathway (Figs. 4 and 6)^42,43^. With reward, SST connectivity reorganizes dramatically, favouring PV-mediated disinhibition and increasing V1 receptivity to reward-signalling feedback. Previous work in V1 have shown that reward association enhances stimulus-specific V1 representations with learning^42^, inducing selectivity changes in VIP, SST, and PV during discrimination and SST suppression for relevant stimuli^8,43^. We did not observe a net reduction in SST inhibition probably due to the nature of the stimulus: the 20 degree stimulus we used engages less SST inhibition compared to the full screen stimulus^31^ that was used before. Most importantly, these studies focused on population shifts without isolating synaptic loci. Our advance is identifying SST→PV strengthening (∼3-fold) as a gain mechanism^44^ distinct from the locomotion-driven VIP→SST disinhibition^27,28^.

What drives the pronounced reorganization of SST outputs during reward association? We observed an increase in feedback input to SST interneurons under reward conditions (Blue box in Fig. 4B), and top-down external inputs have been shown to produce local synaptic potentiation in cortical circuits^45^. Enhanced feedback drive might therefore induce experience-dependent plasticity in SST interneurons during reward association. Notably, the circuit adjustments we observed differed between SST→PC and SST→PV pathways (Fig. 7), suggesting selective modulation of SST outputs. In pyramidal neurons, synaptic boutons targeting different interneuron classes exhibit distinct structural and functional properties^46,47^, demonstrating that cortical neurons can regulate outputs in a target-specific manner. Our findings rise the possibility that SST interneurons modulate their synapses onto PCs and PV interneurons independently. Alternatively, the enhanced SST→PV influence may arise from plasticity within a specialized subpopulation of SSTs, consistent with recent electron microscopy evidence for heterogeneity among SST projections^45,48,49^.

Despite their distinct circuit loci, the net effect of both forms of learning was to shift the PC population towards the sensitizing form of adaptation (Fig. 1 and 5). In naïve mice, there is a wide distribution of adaptive effects across the PC population but of the two types of inhibitory neurons making direct connections to PCs, PVs are predominantly sensitizing while SSTs are depressing (Fig. 2). Optogenetic manipulations indicate that the form of adaptation expressed in an individual PC reflects the relative strength of these inputs, with a decrease in the PV:SST input ratio shifting the neuron towards sensitization^6^. The PV:SST input ratio also fell in both the habituated and rewarded group of mice (Fig. 4C and D), indicating that changes in the balance between these inhibitory inputs also determined the shift in the direction of fast adaptation during these forms of learning (Fig. 7).

These results provide a mechanistic bridge between fast adaptation and slower learning: long-term plasticity can be understood both as changes in long-range inputs to V1 and as selectively reweighting inhibitory pathways that determine where each neuron sits on the depression–sensitization continuum. Previous studies have linked stimulus familiarity to reduced PV activity and increased SST recruitment in V1, from which the roles of different interneurons have been inferred^8,24,50^. Our model-guided optogenetic approach now provides direct evidence that weakens VIP→SST connections and strengthens SST outputs, directly linking these to sensitization.

## Acknowledgements

This work was supported by BBSRC grant BB/X009386/1, a PhD scholarship to SD from the Leverhulme Trust (DS-2017-011) and a Researchers at Risk Fellowship to YeK from the British Academy and Council for At-Risk Academics (RaR/100503).

## Methods

### Animal details

All experimental procedures were carried out in accordance with the UK Animals Scientific Procedures Act (1986) under the approved personal and project licenses granted by the UK Home Office. Experiments were conducted at the University of Sussex under the corresponding establishment license after review by the University of Sussex Animal Welfare and Ethical Review Board.

The animals were housed in polypropylene cages (21 cm × 35 cm) that contained nesting material, a shelter, a treadmill, a tunnel and wooden sticks. Mice were housed in groups of 2 - 3, except in some instances when reintegration into a group was unsuccessful after surgery. All animals were kept on an inverted 12-hour light-dark cycle, with the dark phase from 9:00 am to 9:00 pm and the light phase from 9:00 pm to 9:00 am. We used a total of 50 transgenic adult mice (C57BL/6J background) of either sex at ages ranging from 90 - 240 days. Transgenic mouse lines used were VIP-Cre (VIP tm1(cre)Zjh/J Jackson #010908), PV-Cre (Pvalb tm1(cre)Arbr/J, Jackson #008069), SST-Cre (SST tm2.1(cre)Zjh/J, Jackson #013044) and SST-Flp (Sst tm3.1(flpo)Zjh; Jackson #028579) x VIP-Cre.

### Surgical procedures

All mice underwent surgical preparation for two-photon imaging: Animals were anesthetized with isoflurane (4% induction, 1–2% maintenance) and secured in a stereotactic frame on a heating pad with feedback control. Mice were injected with the steroid Dexamethasone, as well as Metacam and Buprenorphine for pain relief. Additionally, mice were injected with 500 - 1000 μl of sterile saline to prevent dehydration. Under aseptic conditions, the scalp and periosteum were removed, and the skull was scored with a scalpel. A lightweight titanium head-post was attached to the skull using acrylic dental cement (Unifast Trad). A craniotomy (3 mm diameter) was then performed unilaterally above V1, centred at 2.5 - 2.8 mm mediolateral and 3.5 - 4.0 mm posterior to Bregma. Viral injections (1 - 1.2 μl total volume) were administered at two sites around the centre of the craniotomy using a glass micropipette (Drummond #5-000-1001-X) and a Hamilton syringe (80501) mounted on a stereotactic microinjector (WPI UMP3, UMC4 controller). Injections were delivered at a depth of 250 - 350 μm at a rate of 30 nl/min. The pipette remained in place for 5 minutes post-injection to minimize backflow. To image calcium activity in PCs, mice were injected with AAV1.CaMKII.GCaMP6f.WPRE.SV40. To image interneurons we expressed AAV9.CAG.Flex.GCaMP6f.WPRE.SV40. For optogenetic manipulation we used rAAV9/Syn.Flex.ChrimsonR.tdTomato to activate interneurons and AAV5.CBA.Flex.ArchT-tdTomato.WPRE.SV40 to inhibit them. Following injection, the craniotomy was sealed with a cranial window composed of three round coverslips (two 3 mm and one 5 mm, #1 thickness) bonded with optical glue (NOA61, Norland Products), then fixed using superglue and dental cement. Mice were monitored during recovery and returned to their home cage after regaining mobility. Metacam was provided for 3 days post-surgery for pain relief.

After a minimum of 4 weeks, mice were habituated to handling and head-fixation on a treadmill over 1 - 2 weeks until they learned to run consistently. Imaging sessions started no earlier than 6 weeks post-viral injection.

### *In vivo* multiphoton imaging

Calcium imaging was performed with a Scientifica two-photon microscope (SP1, galvanometer mirrors) using a 16x water-immersion objective (Nikon, NA 0.8). Excitation light was provided by a Ti:sapphire laser (Chameleon II, Coherent), tuned between 915 - 940 nm. Images were acquired with a 1.3 zoom, frame rate of 6.07 Hz and a resolution of 256 × 200 pixels using the software ScanImage 5 (Vidrio Technologies). The resulting fields of view measured 400 × 500 μm. Imaging was done in layer 2/3 of V1, at depths ranging from 150 to 300 μm below the brain surface.

### Visual stimuli

Mice were presented with circular, sinusoidal drifting gratings shown on two gamma-corrected LED backlit monitors (BenQ XL2410T, isoluminant at 25 cd/m2, 120 Hz refresh rate) positioned at 45⁰ to the longitudinal axis of the animals. The gratings were generated by the Python library PsychoPy^51^ at a size of 20⁰ of the visual field, a contrast of 100% and a spatial and temporal frequency of 0.04 cpd and 1 Hz respectively. Before the first imaging session the location of the stimulus on the screen was adjusted for each FOV by empirically testing the ability to activate enough neurons within the FOV.

### Experimental protocol

In each imaging session, animals were exposed to the upwards drifting (270⁰) grating in 10 stimulus trials per FOV. Each stimulus trial consisted of a 20 s inter-trial-interval (ITI) showing a uniform grey screen and a 10 s stimulus presentation. Depending on the GCaMP6 expression levels after viral injection, each mouse was imaged in 2 – 10 FOVs. Mice with a low number of FOVs were exposed to additional sets of stimuli in each session (without imaging), ensuring at least 50 stimulus repetitions per session. Whenever we tested the effects of optogenetic manipulation, the stimulus was shown 20 times per FOV of which every other presentation was paired with LED illumination.

To investigate the long-term changes of cell activity caused by repeated stimulus exposure, animals were exposed to the stimulus in overall 6 sessions at an interval of 2-3 days between sessions. In a 7^th^ control session, we exposed mice to a grating drifting in the orthogonal direction (0⁰) to test if the effects of repeated exposure were stimulus specific and reversable. The same FOVs were tracked repeatedly over the sessions, but individual cells were not matched. All analysis therefore reflects populational level changes within each FOV. Mice were divided into two groups: 1) a habituated group, in which animals were simply exposed to the stimulus as described above, and 2) a rewarded group in which animals were rewarded with a drop of water sweetened with condensed milk (mixed 3:1) at the end of each stimulus presentation. The ITI was randomized to be between 10 – 20 s for rewarded animals to prevent time-dependent expectation of the reward. Furthermore, mice from the rewarded group were water restricted and only given fluids during imaging sessions or via syringe in their home cages (>40ml/kg/day). We ensured that they kept a stable body weight of ∼90% of their original weight.

To control for the effects of water restriction, overall reward excitement and randomised ITI we repeated the PC experiments in 4 mice that were water restricted but did not receive a reward timed to the stimulus as the rewarded mice. Instead, they received it in the set up at random times during breaks between sets of trials.

### Monitoring and analysis of locomotion

Mice were head-fixed on a cylindrical treadmill (made from Styrofoam) where they were able to run at their own speed or rest at any time. The rotation of the treadmill was measured using a rotary encoder and directly fed into ScanImage (imaging software) via an analogue DAQ channel and saved as TIFF files for automatic alignment with the two-photon image acquisition. These files were analysed post-hoc with a custom-written script using the software Igor Pro 8/9 (Wavemetrics) and translated into running speed to separate trials in which animals were locomoting from trials in which mice were at rest. A trial was considered as locomoting when an animal ran at a minimum of 3 cm/s for at least 80% of the 10 s stimulus presentation. We only considered locomoting trials for further analysis in this study.

### Monitoring and analysis of pupil size

We recorded the pupil of each animal’s eye contralateral to the hemisphere that had previously been chosen for viral injection, using an infrared camera (DMK 33G618, The imaging source) fitted with a macro lens (Zoom 7000, Navitar) at a framerate of 7 Hz. Each recording frame was triggered by the ScanImage software via the DAQ and automatically aligned with calcium imaging. For post-hoc analysis the acquired AVI files were loaded into Fiji (ImageJ) and turned into binary images (masks) with only black and white pixels using the “threshold” tool. With this tool we manually set an individual threshold for each session (so pixels would be identified as either black or white) in a way that the pupil (black) was fully included but separated from the rest of the image, surrounded by only white pixels. Images were saved as TIFF files and further analysed with a custom-written script in Igor Pro 8/9 that identified the pupil as ROI and measured its area and position in each frame. Whenever the pupil size could not be determined within a frame, due to blinking or grooming of the mice, it was extrapolated between the previous and posterior frames.

### Reward delivery and monitoring of licks

A metal spout for reward delivery was mounted on a piezo sensor placed within a custom 3D-printed frame (developed by the CWW Research Workshop, Charité Berlin). When the animal’s tongue contacted the spout, the resulting vibrations were detected by the sensor and transmitted to an amplifier which allowed for adjustable sensitivity and gain settings. The amplifier’s digital output was directly integrated into the ScanImage software for automatic synchronization with two-photon imaging.

To prevent licking during stimulus presentation, the spout was initially positioned out of the animal’s reach. A servo motor (Geekservo) moved the spout into place only during reward delivery. This movement began 0.25 seconds before the end of the stimulus, so the spout reached the animal precisely at stimulus offset. Because motor movements were detectable by the piezo sensor, the reward was dispensed with a 0.25-second delay via a solenoid pinch valve (WZ-12021-23, Spex® VapLock™). The animal then had a 1-second window to lick before the spout was retracted. Licks were reliably recorded during the stationary phase of the spout’s movement. Mice typically did not miss licking opportunities, as they were introduced to the reward delivery once prior to experiments. This single exposure was generally sufficient to ensure consistent licking behaviour across sessions. Stimulus timing, motor movement, and reward volume were controlled by an ESP32 microcontroller (Espressif Systems) running MicroPython (Beehive).

### Two-photon calcium imaging analysis

The raw imaging data were saved as TIFF files and further processed with MATLAB R2019a (MathWorks) using the Suite-2P package to register the TIFF movies and identify active neurons as regions of interest (ROIs)^52^. All ROIs were manually checked and used for further processing only if they were confirmed to be individual neurons. The extracted calcium traces from these cells were then analysed further with Igor Pro 8 (Wavemetrics), using the SARFIA analysis package^53^ and additional custom-written scripts. To correct for background fluorescence, the average signal (F) of each ROI was adjusted by calculating the mean signal from the surrounding pixels, up to ∼1.5 times the ROI width and excluding overlapping ROIs. The relative intensity of this surrounding signal was then reduced by a “contamination ratio” estimated here at 0.5 and subtracted from the raw signal. The calcium activity of individual cells was expressed as a relative change in fluorescence over time. The baseline fluorescence (F0) for each ROI was determined by identifying the mode of the full calcium trace whenever it was near the minimum. The fluorescence change (ΔF) at each time point was then divided by this baseline (ΔF/F0).

Firing rates were estimated from the recorded ΔF/F data with the MLSpike algorithm^18^. We used parameters expected for GCaMP6f, setting a range of unitary Ca^2+^ transient amplitude between 1 and 50% and the decay time-constant of that transient from 0.05 to 3 seconds. The parameter A_max_, which corresponds to the maximum amplitude of detected events, was 9, and the saturation parameter was set to 0.002. Finally, the Hill coefficient for Ca^2+^ binding to GCaMP6f was varied in the range from 1 to 3.75 with step sizes of 0.25. These parameters were obtained by maximum-likelihood autocalibration of ΔF/F traces, allowing conversion of relative fluorescence changes into inferred spike counts and corresponding firing rates.

Using the StatsLinearCorrelationTest function in Igor Pro, we computed Pearson’s correlation coefficients between each neuron’s ΔF/F activity trace and the stimulus trace binarized as stimulus on (1) or stimulus off (0). Based on this analysis, neurons were categorized as significantly positively correlated, significantly negatively correlated, or not significantly correlated with the stimulus. A p-value threshold of 0.05 was used to determine statistical significance of positively and negatively correlated cells.

To quantify how neuronal responses adapted over the stimulus time, an adaptive index (AI) was calculated for each neuron. This index described the normalized ratio between the early and late response amplitudes during the 10-second stimulus presentation. The initial amplitude was defined as the average ΔF/F measured during the first two seconds of the stimulus (excluding the initial 1 second after stimulus onset), while the final amplitude was taken as the average during the last two seconds. Mean AI values for each condition were computed by averaging across all trials. Cell responsivity was quantified from calcium ΔF/F signals to maximize the number of cells detected across sessions, as the binary classification of responsive versus non-responsive cells is less sensitive to calcium indicator kinetics than precise spike rate dynamics.

### Optogenetic manipulations

Red-shifted optogenetic activators (ChrimsonR or ArchT) were excited through the objective using an amber LED (Thorlabs, 590 nm, M590l3) controlled through a high-power LED driver (DC2200). To prevent contamination of GCaMP signals, the LED was pulsed to deliver light only during the turning phase of the x-mirror of the microscope, as monitored through the position signal of the mirror controller (Cambridge Technology). An Arduino read the position signal from the mirror controller and delivered a TTL signal for the LED driver. Under our usual imaging conditions, the LED power was pulsed at 2 KHz, each pulse lasting ∼0.1 ms.

Where the effect of the optogenetic manipulation was to inhibit PCs, it was important not to completely suppress the initial response so as to quantify a change in AI. LED power was therefore calibrated during each imaging session to ensure that the average initial response of a population of PCs within a FOV was not reduced by more than 75% or by less than 25%. This was achieved using powers of 8–60 µW out of the objective when using ChrimsonR (equivalent to peak intensities of 20–150 µW mm^−2^ at the focal plane).Where the effect of the optogenetic manipulation was to increase PC activity, LED power was adjusted to ensure that the initial response of most PCs was increased by at least 50% but not more than ∼100%. This was achieved using powers of 0.6–4.9 mW using ArchT.

To classify PCs based on their response amplitude changes during SST activation, we compared the baseline amplitudes from control trials to those during SST stimulation using a t-test. Cells with statistically significant differences were further divided into two groups: those with increased and those with decreased response amplitudes, depending on the direction of change. This classification was also based on calcium ΔF/F signals to maximize sensitivity.

### Model

We used a rate-based data-driven population model of the network circuit in mouse visual cortex layer 2/3^15^ (code, parameter values and example data sets are available at https://github.com/lagnadoLab/CortexModel-Learning). A system of differential equations was fitted to experimental traces from the four main neuron populations, dividing VIPs in two subpopulations according to their response to the stimulus (PC, PV, SST, and VIP+ and VIP-). The strength of each connection within layer 2/3 was represented by connection weights (w_ji_). Three external inputs were also considered: feedforward input (FF) coming from layer 4 ^54,55^, feedback (FB) from higher order cortices^56,57^ and a slow modulatory input (SS) that is modulated by locomotion and was necessary to fit the dynamics of signals in session 1^15,26^.

To obtain values of connection weights we fit the model simultaneously to the average activity traces of the five neuronal populations using the *LMFIT* library in Python^58^ by minimization of reduced chi-square parameter (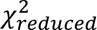) starting from a set of initial conditions. Due to the high dimensionality of the parameter space, multiple initial conditions produced a set of weights that gave acceptable fits with similar 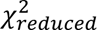 values. To search for the population of acceptable fits, we sampled the whole parameter space using ∼100.000 initial conditions from a combination of homogeneous Sobol sampling^59^ and manual initialization to select the sets of connection weights with lower 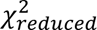 thresholds were determined by inspecting all the tested initial conditions and selecting a cutoff corresponding to fits that showed good agreement with the experimental activity traces. Unsupervised clustering analysis identified a single cluster of acceptable solutions for both conditions in session1 and session 6, indicating that these parameter sets form a continuum rather than falling into distinct categories. The average is, therefore, a representative value of the whole population of acceptable solutions and was used to make further interpretations (plotted in Fig. 3 and 4).

Most of the rest of parameters were fixed based on experimental measurements from the literature or our own recordings^15^. To capture condition-dependent effects while reducing the degrees of freedom of the model, external input parameters were fixed separately for habituated and rewarded traces but were held constant across sessions within each condition. This constraint allowed us to capture differences in dynamics between conditions while focusing the changes on effective connection strengths.

The total synaptic input that neuron i receives from j, Iji, is proportional to the activity of j and can be expressed as:

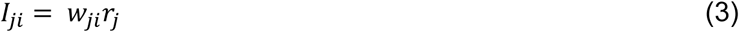

### Statistics

For all statistical tests we used the analytical software Igor Pro 8 and/or SPSS (’Statistical Package for the Social Sciences’, IBM). Data sets were tested for normality with a Kolmogorov-Smirnov (KS) test to determine whether a parametric or a non-parametric test should be used to test for significant differences.

Whenever data was compared between sessions of repeated stimulus exposure, we used linear mixed modelling (LMM) to test for significant differences between session 1 and session 6 within one condition (habituated or rewarded) and to test for significant interaction between conditions and sessions (sess.*cond.). Session and condition were thereby defined as fixed variables and the animal ID was defined as the random variable. The interaction between session and condition was used to detect differential changes between habituation and reward association. We additionally tested the same data within one condition for significant differences between sessions with a Kruskal-Wallis test (KW) with similar results. To assess statistical difference between group of mice in locomotion and pupil size we used Kruskal-Wallis test. All data shown are given as mean ± standard error of the mean (SEM).

## Supplementary Figures

**Supplementary Figure 1.**
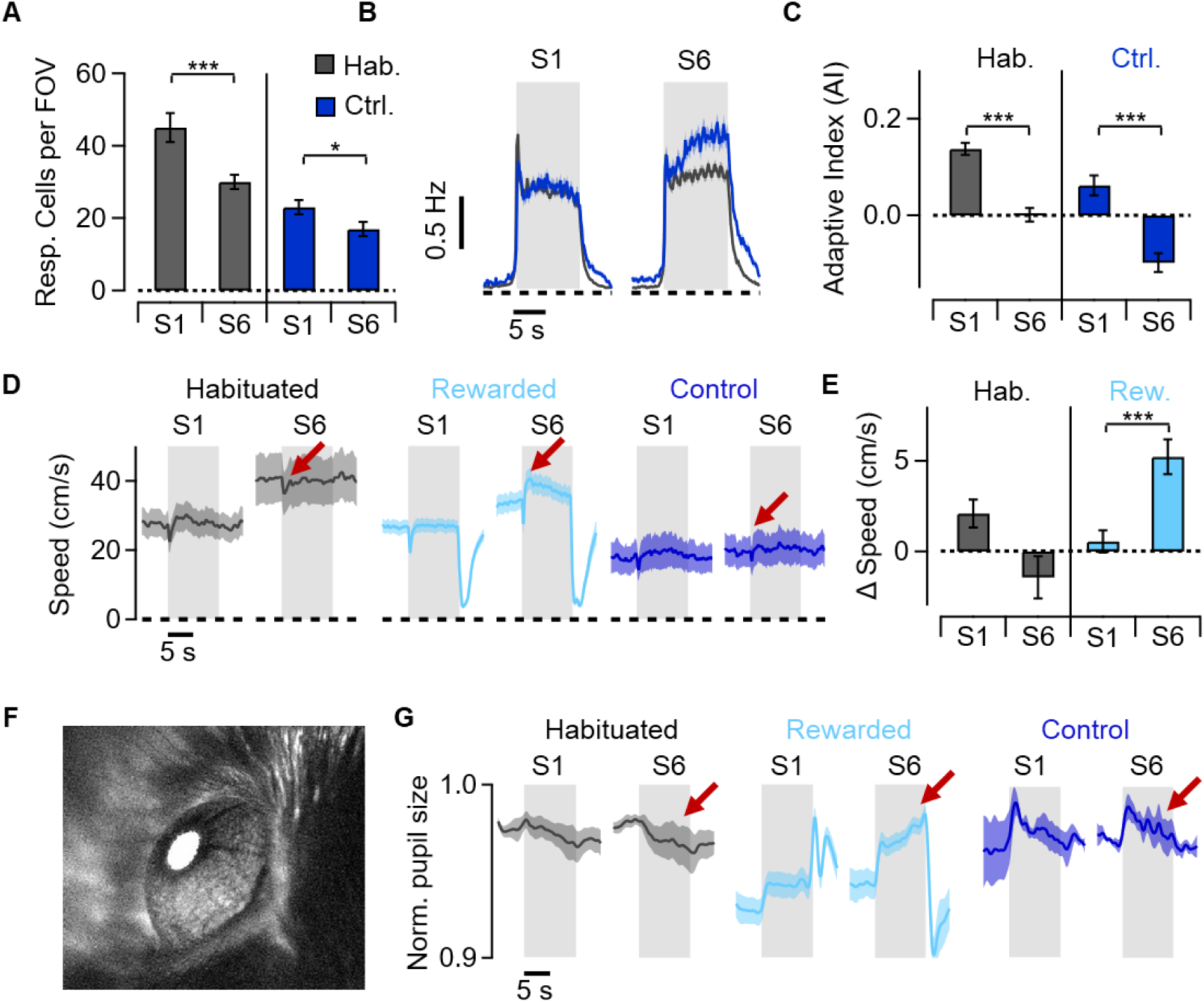
Water-restriction controls and behaviour. Rewarded mice differed from habituated ones beyond stimulus association with the reward: they were water restricted, received a reward in the set up and had variable ITI. To control for these differences, we introduced a control group (n = 4) that was water deprived but received a reward in the setup not associated with the stimulus. **A**. Average number of responsive PCs in habituated and control mice from S1 and S6. During both conditions, the number of responsive cells decreased significantly across sessions (LMM: p < 0.001 for both). The control group had lower overall number of responsive cells from S1 like reward association mice probably due to a change in state with the water reward. **B**. Average PC responses to the stimulus in S1 and S6 of habituated (grey) and control (blue) animals. **C**. The average AI of the PC population of habituated and control mice in S1 and S6. In both conditions, the AI significantly decreased over 6 sessions (LMM: p < 0.001 for both). **D**. Running speed of habituated (n = 14), rewarded (n = 19) and control (n=4) animals in S1 and S6. **E**. Bar graphs showing the change in speed (difference between 2s before and 2s after stimulus onset) after stimulus onset. Habituated mice showed no difference, whereas rewarded mice exhibited an increase in speed (KW: p < 0.001) from S1 to S6, indicating their reward expectation after the onset of the stimulus. **F.** Example mouse pupil during an imaging session. **G**. Normalized average pupil size of habituated (n=14), rewarded (n=19) and control (n=4) animals in S1 and S6. The average pupil response remained relatively unresponsive to the stimulus across all sessions in habituated and control mice, while rewarded animals showed strong increase in pupil size during stimulus presentation once the stimulus had been associated with the reward by S6. This indicates that only reward association animals made an association with the stimulus.

**Supplementary Figure 2.**
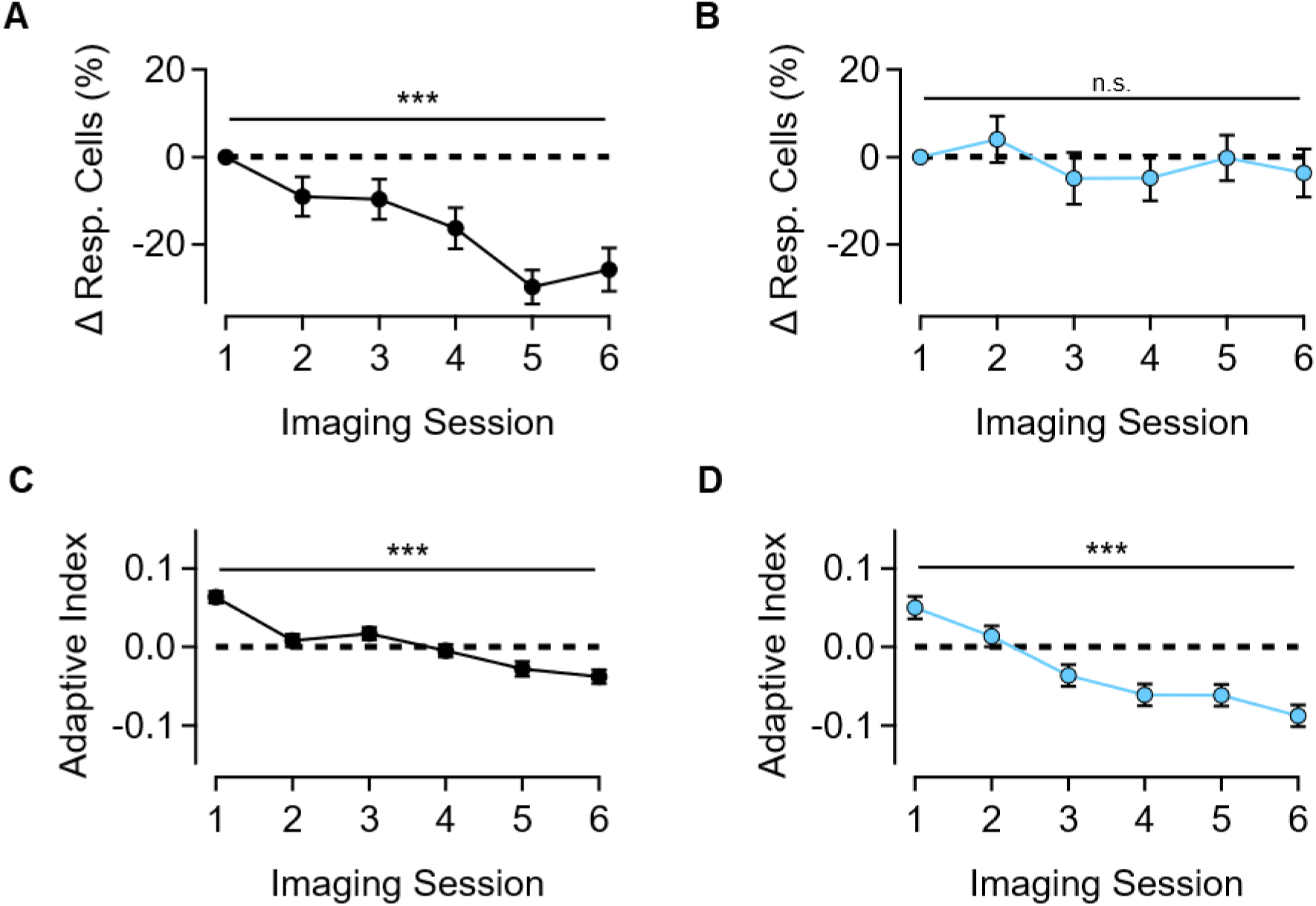
Gradual changes in responsive cells and adaptive index. **A**. The number of responsive cells gradually decreases across 6 sessions in PCs during habituation. (S1 55 FOVs from 14 mice. S1-S6) **B**. same as in A for rewarded animals. Rewarded animals experienced no change in responsive cells across sessions. (S1 30 FOVs from 6 mice. S1-S6. LMM: p = 0.6) **C-D**. A gradual effect was also observed in the shift towards sensitization of PCs during habituation (C. S1 2496 Cells from 14 mice, S6 1633 Cells. S1-S6) and in rewarded animals (D. S1 719 Cells from 6 mice, S6 614 Cells. S1-S6). AI calculated from calcium traces, which was consistent with AI from inferred spikes rates in S1 and S6. Linear mixed model (LMM) ***p < 0.01.

**Supplementary Figure 3.**
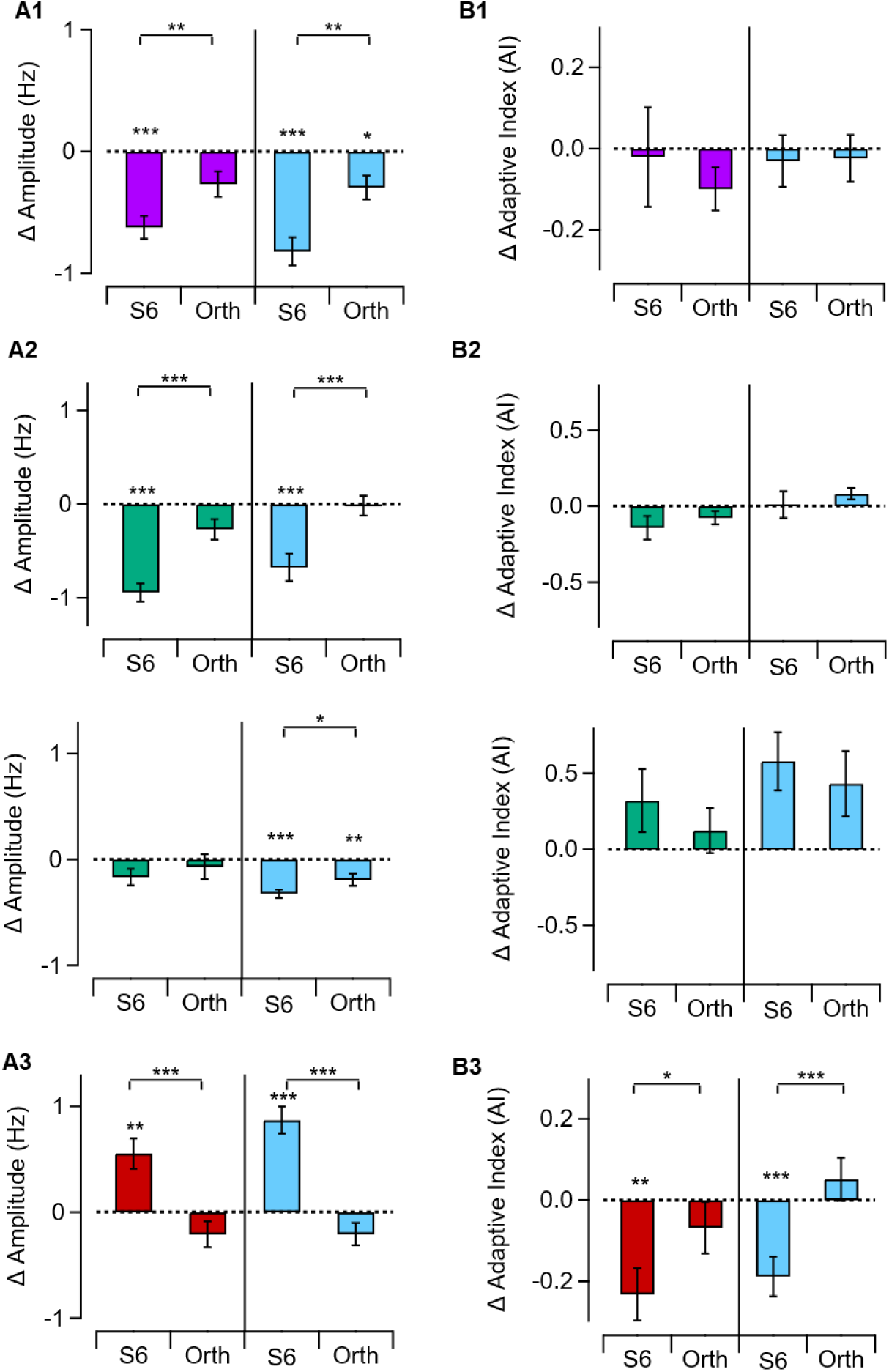
Changes in Amplitude and adaptive index of interneurons A1. Bar plot showing the change in response amplitude in S6 and S7 (orthogonal), both compared to S1, in PV interneurons showing a decrease in both habituated and rewarded mice (Habituation: 19 FOVs from 3 mice. Reward: 22 FOVs from 4 mice). **B1**. Bar plot showing the change in adaptive index in PV interneurons, which remained unchanged. **A2**. Change in response amplitude of VIP+ cells (top) and VIP- cells (bottom). VIP+ decreased in amplitude while VIP- remained unchanged. (VIP+. Habituation: 20 FOVs from 3 mice. Rewarded: 26 FOVs from 4 mice) (VIP-. Habituation: 8 FOVs from 3 mice. Rewarded: 5 FOVs from 4 mice). **B2**. Change in AI of VIP+ cells (top) and VIP- cells (bottom). Notice that it did not vary significantly across the sessions. **A3**. Change in response amplitude of SST interneurons. Unlike the other interneuron types, SSTs increased in amplitude in habituated and rewarded mice (Habituation: 25 FOVs from 4 mice. Rewarded: 29 FOVs from 5 mice). **B2**. Change in AI of SST interneurons, shifting towards sensitization like PC cells in both conditions. Linear mixed model (LMM) ***p < 0.01.

**Supplementary Table 1.** Changes in cell responsivity are stimulus specific. **A.** Number of stimulus-responsive neurons in Session 1 and in response to an orthogonal stimulus in session 7. The change in responsivity was not significant across most conditions and neuron types, showing that the change in responsivity from S1 to S6 was stimulus specific. (LMM, *p < 0.05) (Number of mice: PC Hab 14, PC Rew 6, PV Hab 3, PV Rew 4, VIP Hab 3, VIP Rew 4, SST Hab 4, SST Rew 5).

| Neuron type | Number (FOVs) |  | Responsive Cells per FOV<sup>A</sup> |  |  |  |
| --- | --- | --- | --- | --- | --- | --- |
|  | Hab. | Rew. | Hab S1 | Hab S7 | Rew S1 | Rew S7 |
| PC | 55 | 30 | 45 ± 4 | 39 ± 3 | 22 ± 3 | 23 ± 3 |
| PV | 21 | 23 | 30 ± 3 | 22 ± 2* | 33 ± 3 | 30 ± 3 |
| VIP+ | 22 | 29 | 39 ± 3 | 40 ± 3 | 39 ± 3 | 46 ± 3 |
| VIP- | 22 | 29 | 3 ± 1 | 5 ± 1 | 4 ± 1 | 4 ± 1 |
| SST | 25 | 32 | 11 ± 1 | 11 ± 1 | 16 ± 2 | 12 ± 1 |

**Supplementary Figure 4.**
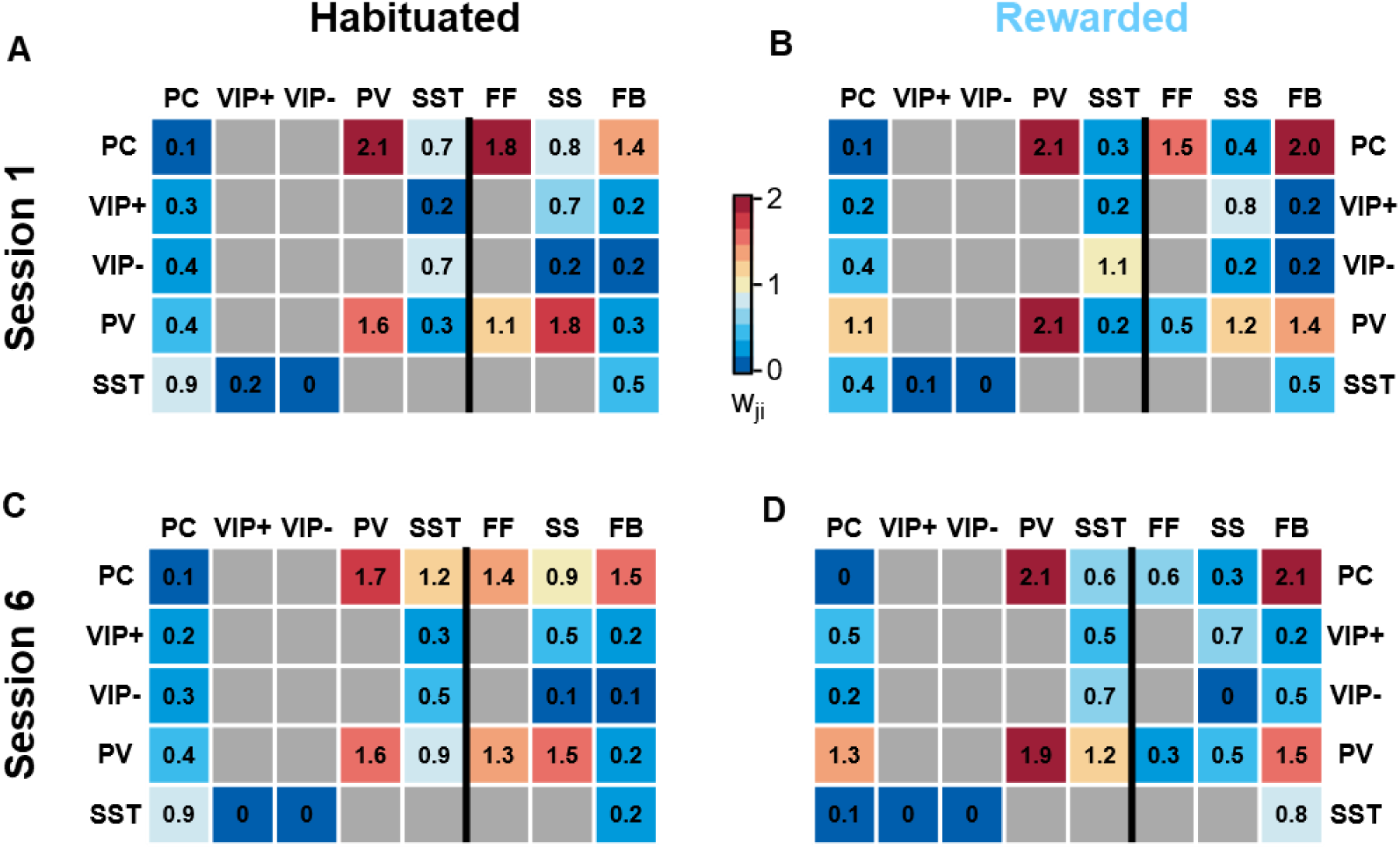
Connection weights of the model. Average connection weights for the set of acceptable solutions in each condition. These were obtained by fitting the model to the activity of responsive neurons in S1 of habituated (**A**; 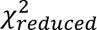< 9, n= 379 solutions) and rewarded mice (**B**; 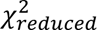< 11, n= 174 solutions), and session 6 of habituated (**C**; 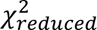 < 6.4, n= 736 solutions) and rewarded mice (**D**; 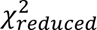< 12, n= 55 solutions). Acceptable solutions were defined as those with reduced chi-square (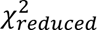) values below a threshold selected for each condition based on their agreement with the experimental activity traces (See Methods).

